# Mechanical loading modulates AMPK and mTOR signaling in myblasts

**DOI:** 10.1101/2024.02.02.578567

**Authors:** Xin Zhou, Shaocun Zhu, Junhong Li, Andre Mateus, Ludvig J. Backman

## Abstract

Skeletal muscle adaptation to exercise involves various phenotypic changes that enhance metabolic and contractile functions. One key regulator of these adaptive responses is the activation of AMPK, influenced by exercise intensity. However, the mechanistic understanding of AMPK activation during exercise remains incomplete. In this study, we utilized an in vitro model to investigate the effects of mechanical loading on AMPK activation and its interplay with the mTOR signaling pathway. Proteomic analysis of myoblasts subjected to static loading (SL) revealed distinct quantitative protein alterations associated with RNA metabolism, with 10% SL inducing the most pronounced response compared to lower intensity of 5% and 2% as well as control. Additionally, 10% SL suppressed RNA and protein synthesis, while activating AMPK and inhibiting the mTOR pathway. Our RNA sequencing analysis further corroborated these findings, revealing numerous differentially regulated genes and signaling pathways influenced by both AMPK and mTOR. Further examination showed that SL induced changes in mitochondrial biogenesis and the ADP/ATP ratio. These findings provide novel insights into the cellular responses to mechanical loading and shed light on the intricate AMPK-mTOR regulatory network in myoblasts.

## Introduction

Skeletal muscle adaptation to exercise involves a multitude of phenotypic changes that contribute to improved metabolic and contractile functions [1]. These adaptations include enhanced mitochondrial quality, increased glucose uptake, and improved insulin sensitivity [1]. An essential player in these adaptive responses is the activation of AMPK (5′-AMP-activated protein kinase) [2]. The activation of AMPK during skeletal muscle contraction is influenced by exercise intensity, with high-intensity exercise resulting in greater AMPK activation compared to low-intensity exercise [3]. AMPK activation is closely tied to the AMP/ATP ratio, which rises due to significant ATP depletion during exercise. Although the mechanisms by which exercise induces AMPK activation remain to be fully elucidated, the use of in vitro models can provide valuable insights into this process.

To mimic the loading patterns experienced by skeletal muscle in vivo, myoblasts can be subjected to mechanical loading using the FlexCell Tension system in vitro. However, direct evidence regarding whether mechanical loading induces AMPK activation in vitro is currently lacking. It is demonstrated that AMPK phosphorylation occur after in situ muscle contraction in rats [4]. In addition, mechanical loading has been shown to activate AMPKγ3 and upregulate the mammalian/mechanistic target of rapamycin, mTOR signaling [5]. Unlike active muscle contraction observed in vivo, the FlexCell Tension system induces elongation of myoblasts to mimic the strain of muscle cells in vivo, i.e. passive strain. The potential of this passive strain to activate AMPK remains unknown. mTOR, is a major regulator of mRNA translation/protein synthesis. It functions in two different complexes, mTORC1 and mTORC2, which regulate cell growth and survival, respectively [6, 7]. Current literature suggests that mTORC1 plays a critical role in stimulating mRNA translation/protein synthesis in skeletal muscle [8]. In skeletal muscle cells, the interplay between two opposing forces, namely mTORC1 and AMPK, governs muscle adaptation to exercise. mTORC1 promotes muscle growth by mediating the anabolic response to resistance exercise, while AMPK is activated during endurance exercises to activate catabolic processes that ultimately lead to normalisation of the AMP/ATP ratio [9]. A computational model suggests that AMPK stimulation subsequently reduces mTORC1 activation [10]. Notably, a connecting aspect of exercise to AMPK activation in skeletal muscle is that exercise induced upregulation of AMPK signaling and downregulation of mTOR signaling [11]. Consequently, exploring the interaction between mTOR and AMPK in myoblasts following various intensities of mechanical loading in vitro is of particular interest.

In this study, we aimed to elucidate the signaling pathways activated in myoblasts subjected to varying static load intensities. Utilizing the FlexCell Tension system, we conducted proteomic analyses, RNA sequencing and supplementary validation experiments. Our results indicate that mechanical loading intensities modulate RNA metabolism in a dose-dependent manner through the interaction between AMPK and mTOR signaling pathways. RNA sequencing further corroborated these findings, revealing several signaling pathways influenced by AMPK and mTOR. Additionally, observed alterations in mitochondrial biogenesis in response to varying loading intensities may underlie the observed AMPK activation.

## Results

### Proteomic Analysis Unveils Disparate Myoblast Responses to Mechanical Stimuli

In order to systematically investigate the cellular alterations induced by mechanical stimuli, we undertook a proteomics analysis of myoblasts. Cells were subjected to three distinct static loading (SL) conditions - 2%, 5%, and 10% - for a duration of 24 hours with intervals of rest in between. Subsequently, cell lysates were obtained for proteomics analysis, resulting in the identification of a total of 6087 proteins. The quantitative outcomes are provided in supplemental data.

When only proteins with a minimum of 2-fold change as compared to the control condition were considered the 2% SL did not induce any quantitative changes. Conversely, in response to 5% SL, 12 proteins were down-regulated, while one protein (ABCC4) was up-regulated. Notably, a more pronounced response was observed in myoblasts following 10% SL, with a total of 68 proteins being affected (Fig 1A). Of particular interest, six proteins exhibited reduced expression levels in both the 5% and 10% SL groups (Fig 1B). Detailed information regarding these proteins, namely RPSA, SUB1/PC4, SUPT5H, SRSF2, RPS21, and PPIL4, is listed in suplementary Table 1. Interestingly, these identified proteins play crucial roles in various steps of the gene expression pathway. Specifically, SUB1 and SUPT5H are involved in promoting RNA transcription and elongation, while SRSF2 is necessary for pre-mRNA splicing. RPSA and RPS21 function as core components of the 40S ribosomal subunit, facilitating mRNA scanning and initiation of protein synthesis. Lastly, PPIL4 accelerates protein folding processes. Pathway and Process Enrichment Analysis revealed that the enriched terms associated with 5% and 10% SL predominantly converged on the metabolism of RNA (Fig 1C).

**Fig 1.**
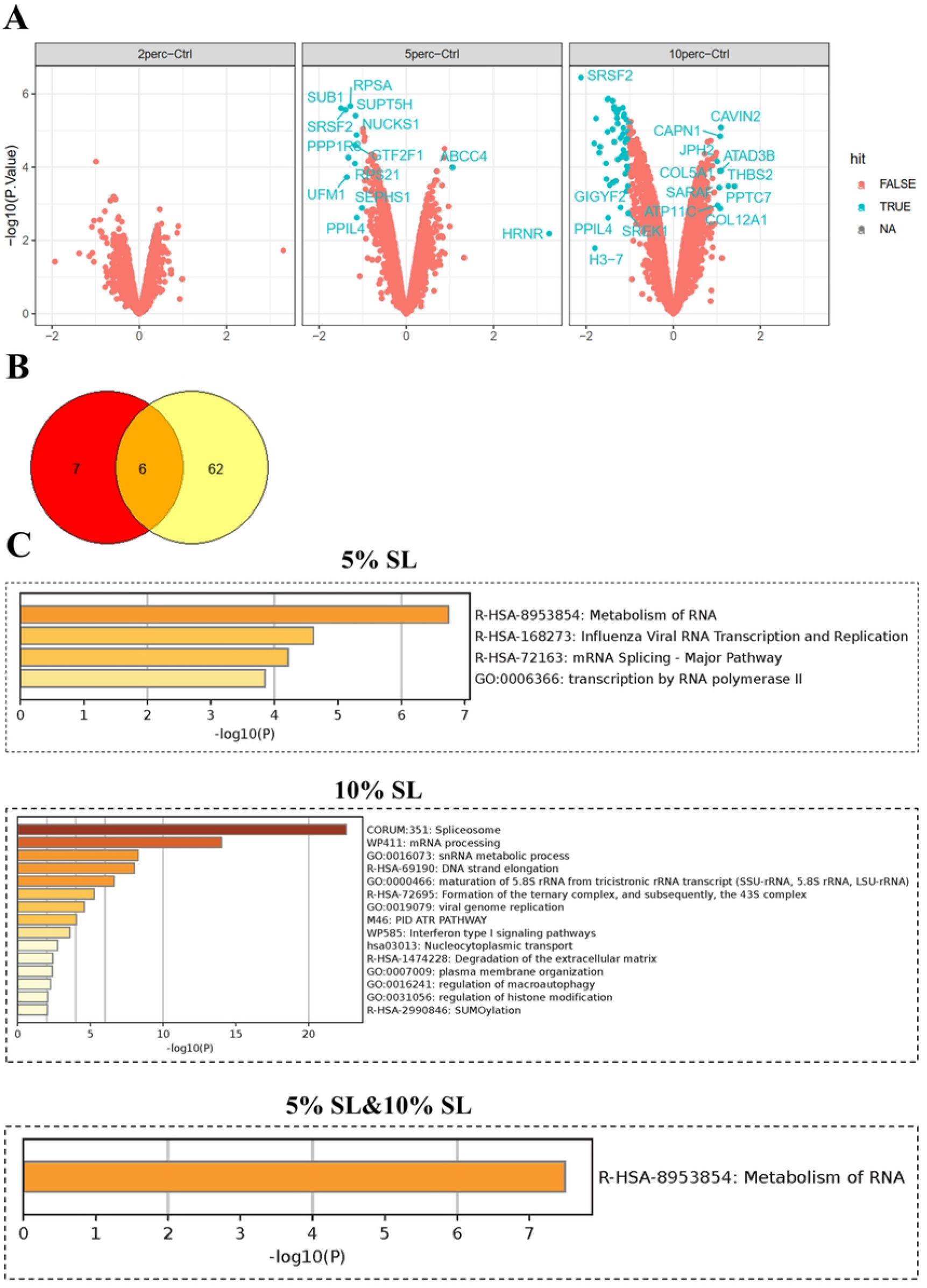
Impact of staticl loading (SL) on RNA Metabolism in myoblasts. (A) The volcano plot presents the differential expression of proteins between the Control and SL groups of 2%, 5%, and 10% for 24 hours. (B) The Venn diagram depicts the overlap of protein identifications between the 5% and 10% SL groups, highlighting shared protein alterations. (C) Pathway and process enrichment analysis using Metascape reveals enrichment of signaling pathways in 5% and 10% SL. Notably, there is convergence in the signaling pathways associated with RNA metabolism between the 5% and 10% SL.

### Statical Loading of 5% and 10% Induces Inhibition of RNA and Protein Synthesis

Given the involvement of the identified proteins in RNA metabolism and protein synthesis, we proceeded to investigate whether these alterations led to a disruption in RNA and protein synthesis. To address this, we employed the Click-iT kit to assess RNA and protein synthesis in response to SL. While 2% SL resulted in an increase in RNA synthesis as compared to control, both 5% and 10% SL led to a significant reduction in RNA synthesis (Fig 2A). Similarly, protein synthesis was prominently reduced in myoblasts subjected to 10% SL as compared to both 2% and 5% (Fig 2B).

**Fig 2.**
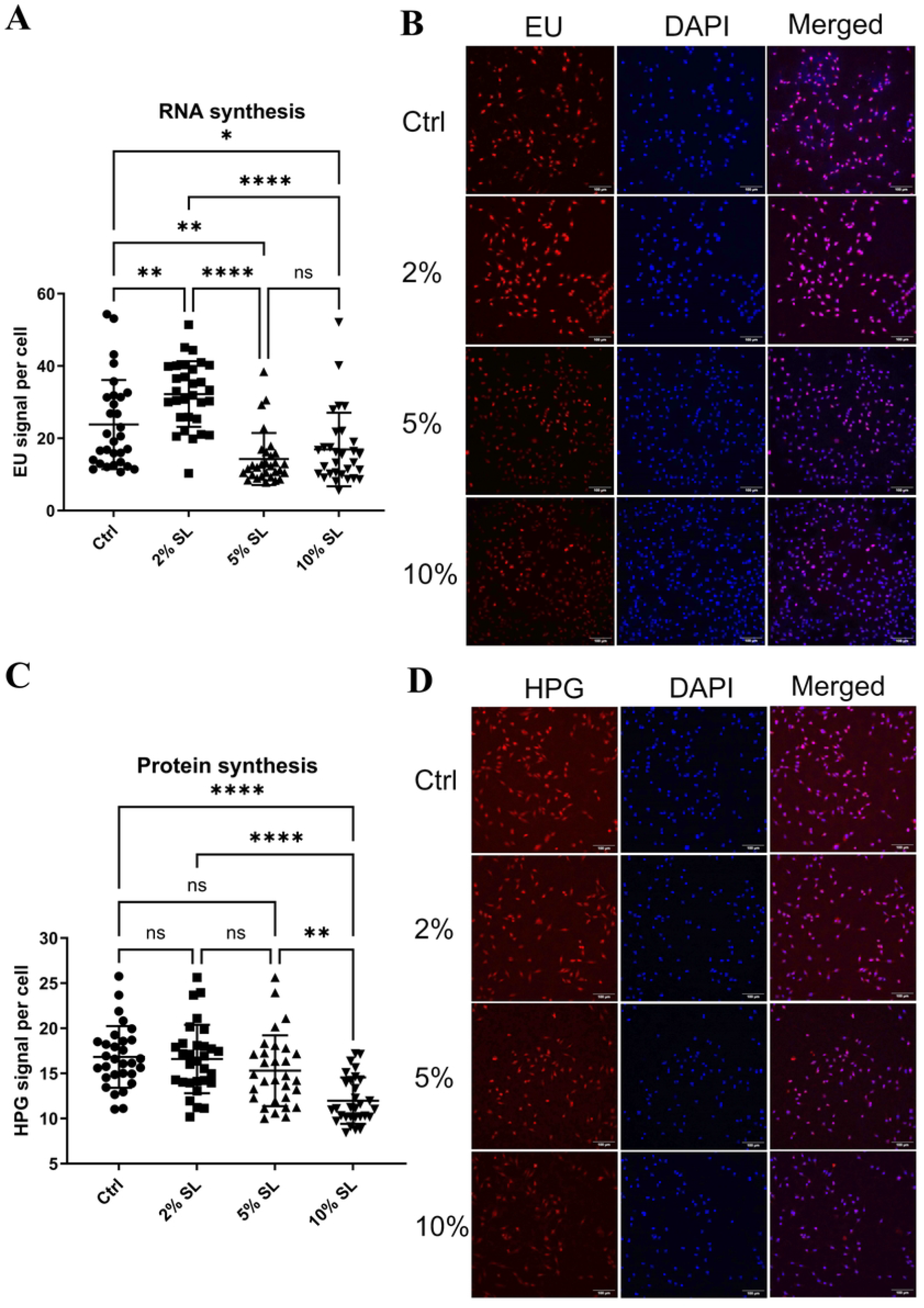
Reduced RNA and Protein Synthesis by 5% and 10% static loading (SL). (A) Visualization and quantification of RNA synthesis in myoblasts following SL using the Click-iT imaging kit. ===== Representative foci images are displayed on the left panel, while the corresponding quantitative data are presented on the right panel. (B) Visualization and quantification of protein synthesis in myoblasts following SL using the Click-iT imaging kit. Representative foci images are presented on the left panel, and the quantitive data are shown on the right panel. The data are presented as mean ± standard deviation. Statistical significance is indicated as ^*^ p < 0.05, ^**^ p < 0.01, ^***^ p < 0.001, ^****^ p < 0.0001.

### 10% Statical Loading Suppresses mTOR Pathway via AMPK Activation

The top 10 significantly altered proteins following 5% and 10% SL are provided in Suplementary Table 2 (Supplemental data). Notably, both SUB1 and SRSF2 are ranked as the top two proteins in 5% and 10% SL groups. Although we encountered difficulties in obtaining an antibody for SUB1 blotting, we present evidence that the expression of SRSF2 is reduced following SL in a dose-dependent manner (Fig 3A), suggesting its potential as a marker for myoblast response to SL. The mTOR pathway, known to regulate cell growth and metabolism in response to mechanical loading [13], plays a critical role in ribosome biogenesis and protein synthesis regulation [12]. Therefore, it is reasonable to investigate whether SL affects mTOR signaling.

**Fig 3.**
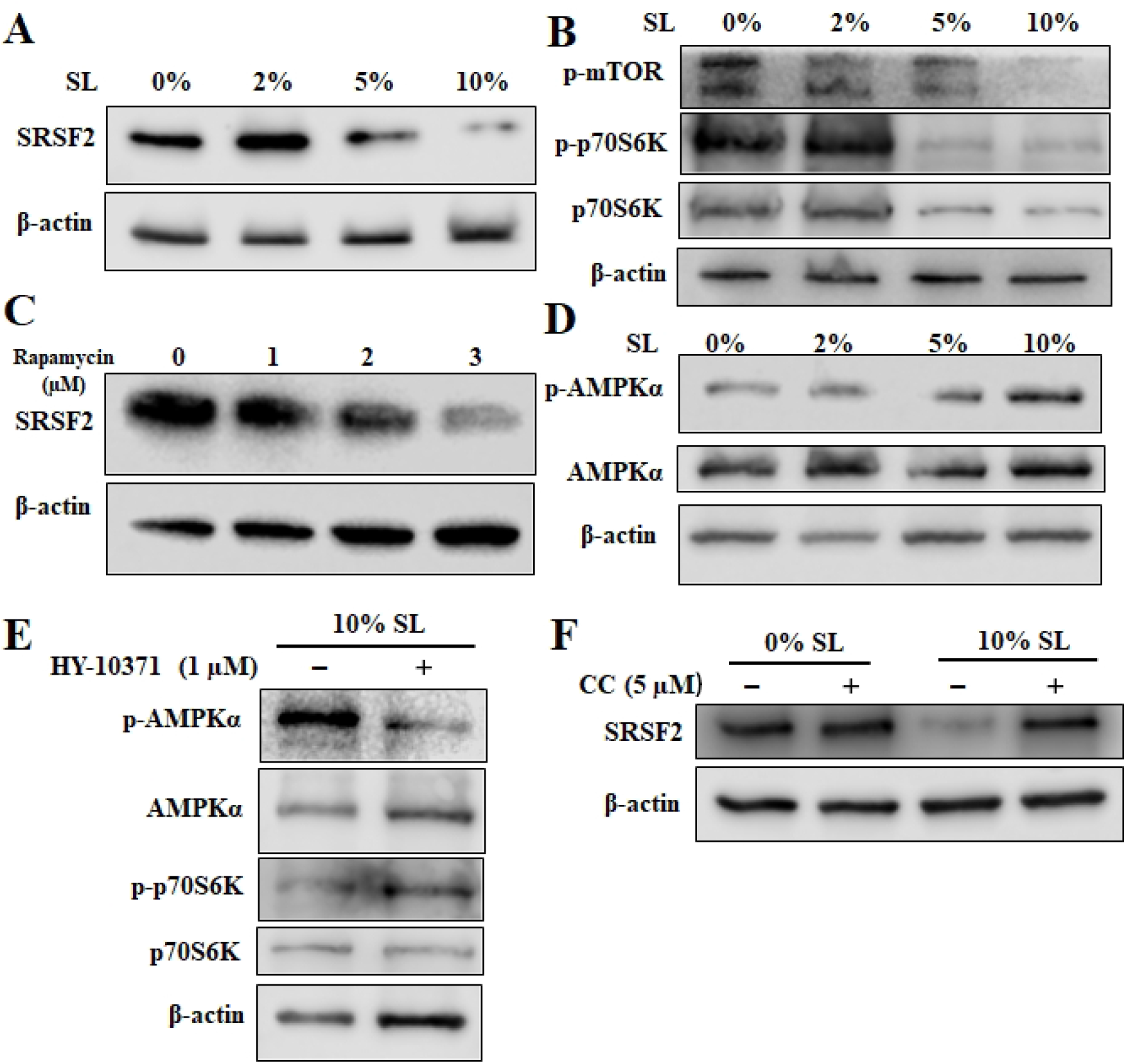
Statical Loading (SL) suppresses mTOR Pathway via AMPK Activation. (A) Decreased expression of SRSF2 in myoblasts in response to increased intensity of SL. (B) Reduced expression of p-mTOR (Ser2448) and p-p70S6K (Ser371) with increased intensity of SL. (C) Myoblasts treated with 1-3 µM rapamycin for 24 hours. The results exhibit a dose-dependent reduction in SRSF2 expression. (D) Increased expression of pAMPKα (Thr172) in myoblasts following SL. (E) HY-10371 pretreatment abolished AMPK phosphorylation and rescued p70S6K phosphorylation in loaded myoblasts. (F) Effect of 5 µM Compound C (CC) treatment on SRSF2 expression in myoblasts exposed to 10% SL. Myoblasts were incubated with or without CC for 24 hours. The addition of CC rescued the expression of SRSF2 in myoblasts subjected to 10% SL. Actin served as a loading control.

The expression of p-mTOR exhibited an intensity-dependent pattern, with reduced levels observed in 2% and 5%, compared to the unloaded control, and diminished expression in 10%. Similar expression pattern was observed for phosphorylation of the protein S6 kinase (p70S6K), a key indicator of mTOR activation. Interestingly, treatment with rapamycin, a known mTOR inhibitor, dose-dependently reduced SRSF2 expression (Fig 3C). These findings confirm that SRSF2 is regulated by mTOR in response to SL.

Additionally, we observed that AMPK signaling was activated in response to increased SL, as evidenced by elevated levels of pAMPKα in myoblasts following 5% and 10% SL (Fig 3D). AMPK activation requires LKB1 phosphorylation. Even though we didn’t find suitable antibodies to blot pLKB1, our data showed that AMPK activation induced by 10% SL was markedly reduced by the addition of LKB1 inhibitor HY-10371 (Fig 3E), suggesting the important role of LKB1 in mediating mechanical loading-induced AMPK activation. Furthermore, we observed that treatment with HY-10371 in 10% SL led to a rescue of p70S6K expression. This finding provides additional confirmation of the interplay between AMPK signaling and mTOR pathway (Fig 3E). To further explore the relationship between AMPK and SRSF2 expression, we utilized Compound C, a specific AMPK inhibitor, which rescued the inhibitory effect of SL on SRSF2 expression (Fig 3E). These data suggest that higher intensity of SL inhibits the mTOR pathway through AMPK activation. It is worth noting that AMPK activation has been previously reported to suppress mTORC1 activity [13].

### RNA sequencing confirmed the changes of AMPK and mTOR signaling after statical loading

To corroborate our findings on the interaction between AMPK and mTOR pathways, we mapped the mRNA expression profile of statically loaded myoblasts using RNA sequencing. We observed dose-dependent alterations in both AMPK and mTOR pathways, as evidenced by KEGG (Kyoto Encyclopedia of Genes and Genomes) pathway enrichment analysis. Specifically, after 2% SL, AMPK signaling axes indicated enhanced biosynthesis and cell proliferation, marked by upregulated expression of Cyclin D1 and Cyclin A (associated with cell growth), HMGR (cholesterol biosynthesis), SCD1 (unsaturated fatty acid synthesis), 4EBP1 (cell growth and protein biosynthesis), and downregulated Ulk1 (autophagy) (Fig 4A left panel). Conversely, 10% SL conditions resulted in inverse expression patterns for these markers (Fig 4C left panel). Intermediate 5% SL mirrored the 10% loading trend, displaying reduced expression of Cyclin A, HMGR, and SCD1 (Fig 4B, left panel). Similarly, mTOR signaling profiles exhibited dose-dependent variations. Notably, PRAS40 and Deptor, two key inhibitors of mTOR signaling, were downregulated in the 5% and 10% SL groups, while 2% SL did not affect their expression (see Fig 4A, B, C, right panel).

**Fig 4.**
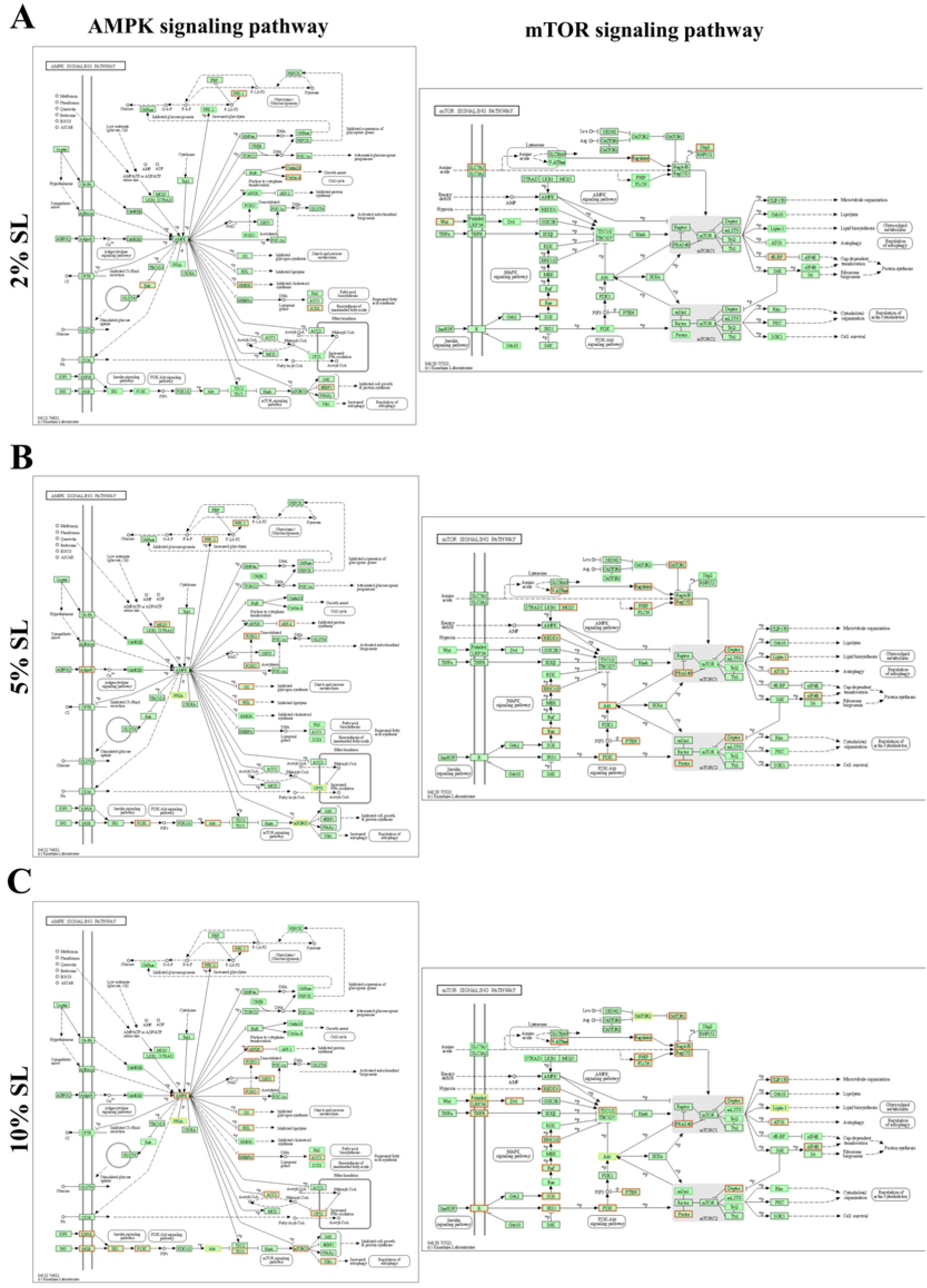
Static loading (SL) induced changes in mRNA profile related to AMPK and mTOR signaling. KEGG Signaling Pathway Enrichment Analysis of mTOR Signaling in Response to SL Treatment. (A-C left panel) show AMPK signaling pathway in response to 2%, 5% and 10% SL. A clear dose-dependent increase in AMPK pathway activation with increase intensity of SL. (A-C right panel) show mTOR signaling pathway in response to 2%, 5% and 10% SL. 2% SL induced activation of the mTOR signaling pathway while 5% and 10% SL exhibited a dose-dependent inhibition of the mTOR signaling pathway. Gene expressions in the red frame are upregulated, in the green frame are downregulated, and those in the yellow frame show variable regulation, either upregulated or downregulated.

### Statical Loading induced elevated mtRNA expression and transit changes in ADP/ATP ratio, mitochondrial membrane potential

AMPK serves as a crucial sensor of cellular energy status, becoming activated in response to elevated AMP and ADP levels. To investigate the dynamics of AMPK activation in myoblasts following 10% SL, we monitored the time course of ADP/ATP ratio. Notably, a rapid increase in ADP/ATP ratio was observed 1 hour after applying 10% SL, which subsequently returned to basal levels at 9 and 24 hours (Fig 5A). Considering the pivotal role of mitochondria in ATP generation, we examined the parallel changes in mitochondrial membrane potential. Intriguingly, we detected significant changes in mitochondrial membrane potential (MMP) 1 hour after 10% SL, which exhibited a gradual recovery over time (Fig 5B), as measured by TMRM staining. Given that mitochondrial positioning and morphology are known to be regulated by the cytoskeleton [14], we employed real-time live imaging to assess the morphological changes following 1 hour of 10% SL. We observed immediate cellular morphology shrinkage, which subsequently returned to normal within 6 hours, in concurrence with MMP (Supplemental data). The intricate relationship between cytoskeleton and energy metabolism prompted us to investigate the potential link between mechanical loading-induced changes in the MMP and the observed shift in ADP/ATP ratio. Notably, our findings suggest that alterations in the MMP following SL contribute to the activation of AMPK signaling.

**Fig 5.**
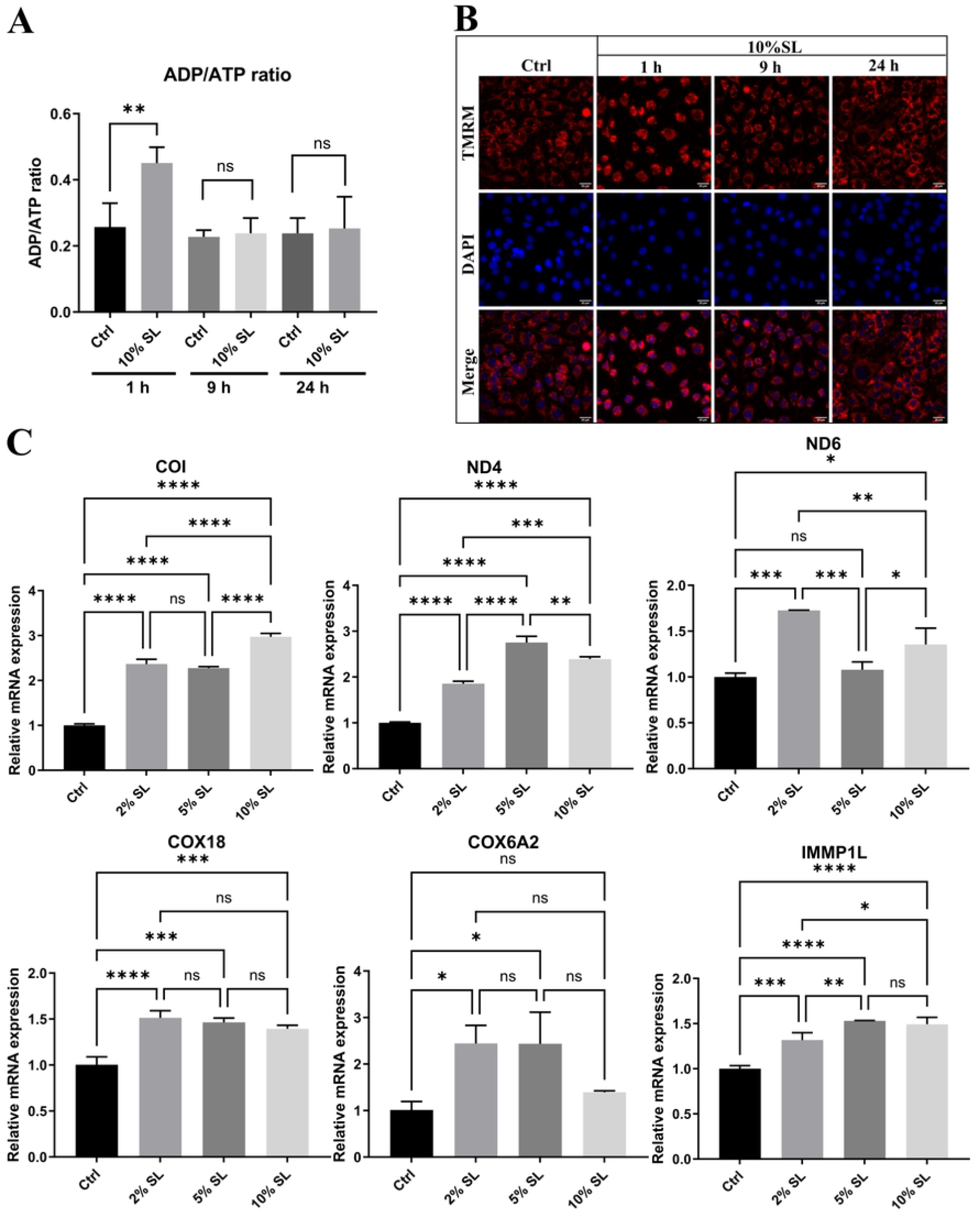
Statical loading (SL) induced changes in parameters related to mitochondrial biogenesis. (A) ADP/ATP ratio measured at 1, 9, 24 hours following 10% SL. Significant increase of ADP/ATP ratio were seen at 1 hour after 10% SL. (B) Mitochondrial membrane potential were visulized by TMRM staining. Hoechst stain 33258 was used to label the nucleus. Representative images were shown at each indicated time point. (C) mRNA expression of mitochondrial proteins encoded by mtDNA (upper panel) and nuclear DNA (lower panel) 24 hours after 10% SL. The data are presented as mean ± standard deviation. Statistical significance is indicated as ^*^ p < 0.05, ^**^ p < 0.01, ^***^ p < 0.001, ^****^ p < 0.0001.

Given the retrospective role of AMPK in stimulating mitochondrial biogenesis, we examined the transcriptional activity of six key genes involved in the electron transport chain (ETC) 24 hours after SL (Fig 5C). Quantitative PCR (qPCR) analysis revealed an overall upregulation of mitochondrial DNA (mtDNA)-encoded genes after SL, including COI, ND6, and ND4, as well as nuclear DNA (nDNA)-encoded genes, including COX18, NDUFAF2, and COX6A2. Among the tested genes, MT-ND4, MT-ND6, and NDUFAF2 are critical components of NADH: Ubiquinone Oxidoreductase (mitochondrial complex I), while COX18, COX6A2, and COI are essential components of Cytochrome C oxidase (mitochondrial complex IV). Notably, a general increase in mtRNA expression was observed for all six genes tested, except for ND6 after 5% SL and COX6A2 after 10% SL. These results strongly suggest the activation of mitochondrial biogenesis, potentially mediated by AMPK signaling activation.

## Discussion

To our knowledge, no study has investigated the proteomic changes in myoblasts after in vitro mechanical loading. The observed intensity-dependent responses of myoblasts to mechanical loading substantiates the reliability of the FlexCell system as a tool for investigating mechanical responses in vitro. Our proteomic profiling offers valuable insights into the distinct responses of myoblasts to different intensities of mechanical stimuli, emphasizing the significance of the AMPK-mTOR signaling pathway. Noteworthy, our proteomics analysis identified four proteins, namely SUB1 (PC4), SRSF2, RPSA, and RPS21, that have been previously associated with mTOR signaling [12]. SUB1 is a upstream regulator of mTOR, as it inhibits the deacetylation activity of Sin3-HDAC, thereby activating mTOR signaling [15]. Unfortunately, suitable antibodies for detecting SUB1 in rat myoblasts were unavailable, necessitating further investigation in future studies. Given that the mTOR pathway governs the synthesis of ribosomal components, including ribosomal proteins (RPs) [12], such as RPSA and RPS21, it is plausible that these proteins are regulated by mTOR. In-depth investigations are required to establish a direct connection between mTOR signaling and RPs. We further showed that SRSF2 expression followed the patten of mTOR activation, and were reversely regulated by AMPK pathway. These findings suggest that SRSF2 expression is regulated by AMPK-mTOR signaling axis.

In vivo study using global phosphoproteomic analysis of human skeletal muscle discerningly unveiled a similar pattern in the AMPK-mTOR crosstalk (Fig 2C in [11]). Similar result is reported by Marin et al. when employing Phosphoproteomics in human, rats and mice [16]. In addition to their findings, our findings also underscore the crosstalk between AMPK and mTOR signaling. Additionally, activation of AMPK was accompanied by dynamic alterations in MMP and upregulation of pivotal genes involved in mitochondrial biogenesis. It is noteworthy that, despite an overall inhibition of RNA synthesis following 10% SL, most expressions of mtRNA were augmented after SL. This outcome strongly supports the notion of enhanced mitochondrial biogenesis after SL. Together with the observed cellular morphological changes and mitochondrial membrane potential, our findings suggest adaptive responses in mitochondria, working in conjunction with the cytoskeleton, to the mechanical loading. These results provide valuable insights into the adaptive responses of myoblasts to mechanical stimuli and their implications for cellular energy metabolism.

In conjunction with our proteomic findings, RNA sequencing data revealed that numerous downstream targets of AMPK, including Cyclin D1, Cyclin A, HMGR, SCD1, 4EBP1, and Ulk1, exhibited loading intensity-dependent transcriptional regulation, aligning with AMPK activation. A similar pattern was observed in the mTOR signaling pathway, where increased expression of mTOR inhibitors PRAS40 and Deptor was noted specifically at 5% and 10% SL. This RNA sequencing not only corroborated the changes in AMPK and mTOR signaling but also provided insights into the broader signaling alterations under mechanical loading.

Overall, our study presents novel findings on the proteomic changes and signaling pathways involved in myoblasts in responses to mechanical loading in vitro. This study contributes to the understanding of the molecular mechanisms underlying muscle adaptation to exercise and highlights the interplay between AMPK and mTOR signaling in this process.

## Materials and Methods

### Cell culture

L6 myoblasts were cultured in T-175 flasks (Sarstedt, #83.3912.002) in Dulbecco’s modified Eagle medium (DMEM; Thermo Fisher Scientific, #31966021) containing L-alanyl-L-glutamine (GlutaMAX) and 10% fetal bovine serum (FBS; Thermo Fisher Scientific, #10500064). Prior to experiments FBS concentration was reduced to 1%.

### Mechanical loading and drug treatments

The FlexCell Tension System (Flexcell International Corporation, FX5000, USA) was used to generate mechanical loading to the adherent myoblasts. FlexCell plates were placed on 25 mm diameter round equibiaxial loading posts. The flexible-bottomed membrane of the FlexCell plates were stretched by vacuum suction. The myoblasts received static mechanical loading (SL) of 2%, 5% and 10%. The loading protocol was as follow: 1 hour SL followed by 2 hours rest period; this SL and rest period was repeated three times before a resting period of 6 hours. The protocol was repeated for 24 hours. By adding rest intervals and applying mechanical strain for intervals, stimulus adaptation is avoided and the mechanical sensitivity of cells maintained [17]. No significant alterations in cell death were observed following the loadings, as determined by lactate dehydrogenase (LDH) assay, data not shown.

For rapamycin treatment, cells were cultured with 1-3 µM rapamycin for 24 hours. For Compound C treatment, cells were cultured with 5 µM Compound C for 24 hours with/without SL. For HY-10371 treatment, cells were cultured with/without 1 µM HY-10371 for 24 hours with 10% SL.

### Proteomics analysis

#### Sample preparation

Medium was switched to SkBM™-2 Basal Medium (SkGM-2 BM, Lanza, #CC-3246) supplemented with SkGM™-2 SingleQuotsTM (SkGM-2 SQ, Lanza, #CC-3244) before loading. Samples were collected and digested into peptides using a modified SP3 protocol [1, 2]. Briefly, cell pellets were resuspended in lysis buffer (2% SDS, 20mM TCEP) and boiled at 95°C for 10 min. SpeedBeads magnetic carboxylate modified particles (Sigma Aldrich, beads A hydrophylic, cat.no GE45152105050250; beads B hydrophobic, cat.no GE65152105050250,) were combined with ratio 1:1 v/v and washed using LC-MS water for four times. Then the beads were mixed with each sample in binding buffer (50% ethanol and 2.5% formic acid in final) and incubated with shaking at 500rpm for 15min at room temperature (RT). Then they were transferred into one filter plate (0.22 µm, Sigma Aldrich, part.no: MSGVN2210). Unbound fraction was removed by centrifugation at 1000rcf. Beads were retained on the filter and washed with 70% ethanol for four time. Trypsin was mixed with digestion solution (100 mM HEPES pH 7.5, 5 mM chloroacetamide, 1.2 mM TCEP) and added to each sample (1 µg trypsin was used for 25 µg protein) on the plate. Samples were digested overnight at RT with shaking at 500rpm. Flowthrough containing peptides was collected with centrifugation at 1000rcf. 10 µl 2% DMSO was added to beads for eluting bound peptides and pooled with the previous flowthrough. Peptides were desalted by the Oasis HLB plate (Waters, cat.no 186001828BA) using the factory protocol and then dried by speed vac.

#### LC-MS/MS

Dried peptides were dissolved with 0.1% formic acid in water. 1µg peptides from each sample was introduced to MS using the Vanquish Neo (Thermo Scientific). The trapping column was PEPMAP NEO C18 (5 µm particle size, 300 µm^*^5mm, Thermo Scientific). Analytical column was nano EaseTM M/Z HSS C18 T3 (100Å, 1.8 µm particle size, 75 µm^*^250mm, Waters). Total length of 2 hours for separation and elution was performed with a gradient of mobile phase A (water and 0.1% formic acid) to 8% B (80% acetonitrile and 0.1% formic acid) over 4 min, and to 27% B over 87 min, then rise to 80% B in 0.1 min and hold for 4 min, finally to 2% B in 30 sec and finally column equilibration was performed.

Data acquisition on Exploris 480 (Thermo Scientific) was carried out using a data dependent method. Survey scans covering the mass range of 375 – 1500 were acquired at a resolution of 120,000, RF lens of 40% and normalized automatic gain control (AGC) of 300%. Maximum cycling time of 2 sec was used to control the number of precursors for tandem-MS/MS (MS2) analysis. Charge states include 2-6 charges. Dynamic exclusion was set to exclude the previously selected precursors for 35 sec. MS2 scans were acquired at a resolution of 15,000 (at m/z 200), with AGC target value of auto. The isolation window was 1.4 m/z. HCD fragmentation was induced with a normalized collision energy (NCE) of 30. Isotopes were excluded for MS2 analysis.

#### Data analysis

Raw data was searched against the homo sapiens UniProt FASTA (proteome identifier [ID] UP000005640) using FragPipe(version 18), label free quantification was achieved using LFQ-MBR workflow. Proteins identified from contaminants and decoyed were removed. Only proteins that quantified in more than one replicate in each group were retained for further analysis. R (version 4.2.2) was used for statistics analysis and volcano plots. To reduce technical variation, data was normalized using vsn package[3]. Protein differential expression was evaluated using the limma package. Differences in protein abundances were statistically determined using the Student’s t-test moderated by Benjamini–Hochberg’s method.

### RNA Extraction and qRT-PCR

Extraction of mRNA was performed using the RNA extraction kit (Qiagen, Venlo, Netherlands, # 74106) according to the manufacturer’s instructions. Subsequently, high-capacity cDNA reverse transcription kit (Thermo Fisher, Waltham, MA, USA) was used to reverse transcribe RNA into cDNA. To determine the gene expression, TaqMan Gene Expression Assays (Applied Biosystems, Carlsbad, CA, USA) were used. cDNA was run using ViiA7 Real-Time PCR system and analyzed with its software (Applied Biosystems, Carlsbad, CA, USA). Gene expression was measured by TaqMan Gene Expression Assay (Applied Biosystems, Carlsbad, CA, USA) and calculated by 2–ΔΔCt method. All probes used for real-time PCR (Applied Biosystems, Carlsbad, CA, USA) are summarized in Table 1. Rpl13a was used as the reference gene for normalization.

**Table 1.**
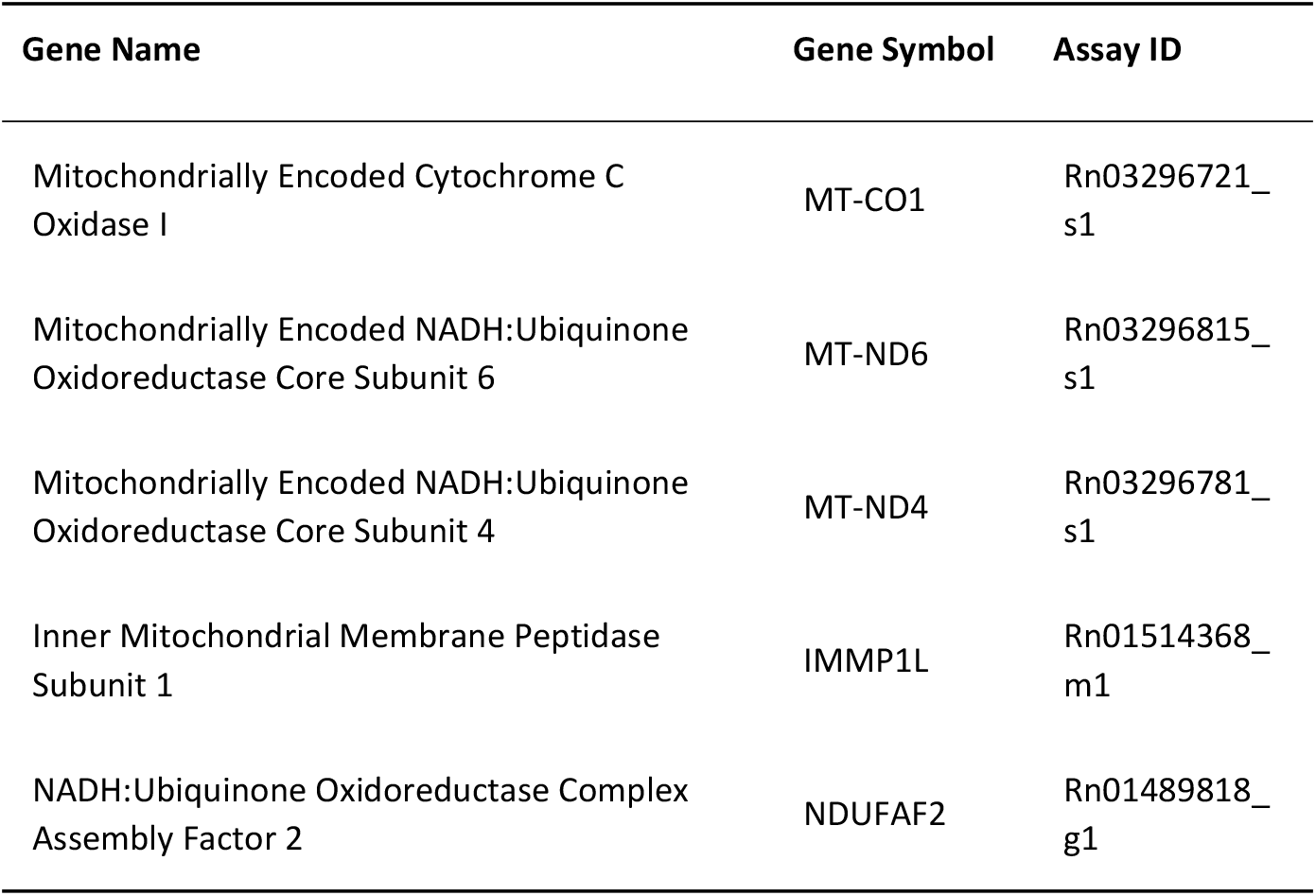
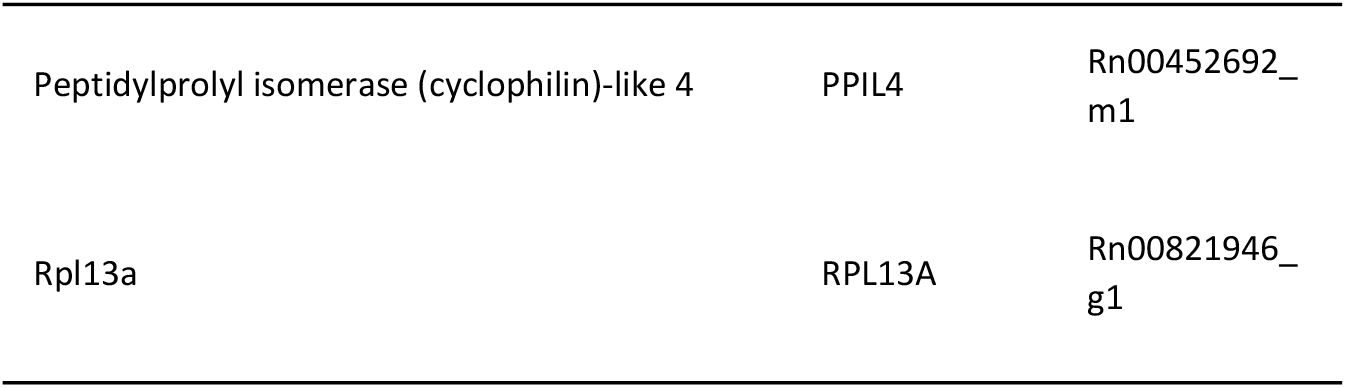
All probes used for real-time PCR.

| Gene Name | Gene Symbol | Assay ID |
| --- | --- | --- |
| Mitochondrially Encoded Cytochrome C Oxidase I | MT-CO1 | Rn03296721_s1 |
| Mitochondrially Encoded NADH:Ubiquinone Oxidoreductase Core Subunit 6 | MT-ND6 | Rn03296815_s1 |
| Mitochondrially Encoded NADH:Ubiquinone Oxidoreductase Core Subunit 4 | MT-ND4 | Rn03296781_s1 |
| Inner Mitochondrial Membrane Peptidase Subunit 1 | IMMP1L | Rn01514368_m1 |
| NADH:Ubiquinone Oxidoreductase Complex Assembly Factor 2 | NDUFAF2 | Rn01489818_g1 |
| Peptidylprolyl isomerase (cyclophilin)-like 4 | PPIL4 | Rn00452692_m1 |
| Rpl13a | RPL13A | Rn00821946_g1 |

### RNA sequencing and bioinformatics

The RNA sequencing process was carried out by Novogene at their Cambridge Genomic Sequencing Centre in the UK, utilizing the Illumina NovaSeq 6000 system. The sequencing commenced with a quality assessment of the samples to ensure they conformed to RNA sequencing standards.

Following this, the library preparation was undertaken, and its quality ascertained. The sequencing approach involved using a 150 bp paired-end strategy for the lncRNA and circRNA libraries, and a 50 bp single-end strategy for the small RNA library. To refine the sequenced data, raw reads were cleansed by removing reads with adapters, those containing more than 10% N bases (where ‘N’ denotes an undetermined base), and reads of low quality (where over 50% of the bases had a Qscore ≤ 5). The HISAT2 tool was employed to align the purified reads to the rat reference genome. The mapping data from all the samples was then processed using the StringTie assembler to assemble the transfrags, which were then compared with reference transcripts to identify potentially novel sequences. Transcript abundance was quantified based on the number of sequenced fragments mapping to exons, using Fragments Per Kilobase of transcript sequence per Millions base pairs sequenced (FPKM) as the measure for gene expression levels. This data was normalized, and the p-values and False Discovery Rate (FDR) were calculated considering multiple hypothesis testing. KEGG analysis was applied to categorize the upregulated and downregulated genes, revealing enriched functions in biological processes and molecular functions for these genes.

### RNA/protein synthesis assay

5-ethynyl uridine (EU) (1 mM final) or L-homopropargylglycine (HPG) (50 µM final) was added to the culture medium for 1 hour to label nascent RNA or protein, and then the cells were fixed and permeabilized as previously described. EU labeling of RNAs was detected using the Click-iT RNA Imaging Kit (Life Technologies, cat. #C10639). HPG labeling of proteins was detected using the Click-iT™ HPG Alexa Fluor™ 594 Protein Synthesis Assay Kit (Life Technologies, cat. #C10429), following the manufacturer’s protocol. The intensity ratio of foci to DAPI was quantified by analyzing ten randomly selected fields from each sample. Three biological repeats were included.

### Western Blot

Cells were freeze-thawed and further lysed in RIPA (radioimmunoprecipitation) lysis buffer (Thermo Fisher, Waltham, MA, USA) supplemented with protease and phosphatase inhibitor cocktail (Sigma, St. Louis, MO, USA, #P1860). Total protein concentration was determined with the BCA assay (Thermo Fisher, Waltham, MA, USA). Samples containing 20 µg of protein were separated on SDS-polyacrylamide gels and transferred to PVDF or NC membranes (Thermo Fisher, Waltham, MA, USA). Membranes were blocked in 5% bovine serum albumin in TBST for 1 hour before staining with primary antibodies overnight at 4°C. After washing, the membranes were stained with HRP-conjugated secondary antibodies for 1 hour at RT before incubation with ECL solution and then analyzed in an Odyssey Fc Dual-Mode Imaging System (LI-COR Biotechnology, Nebraska, USA). β-actin was used to normalize target protein expression. All antibodies used are summarized in Table 2.

**Table 2.** Antibodies used for immunostaining and Western blot.

| Antibody | Company | Code | Applications | Molecular weight (kDa) |
| --- | --- | --- | --- | --- |
| Phospho-AMPK (Thr172) | Cell Signaling | 2535S | WB | 62 |
| AMPKα | Cell Signaling | 2532 | WB | 62 |
| Phospho-mTOR (Ser2448) (D9C2) | Cell Signaling | 5536S | WB | 289 |
| SRSF2 | Thermo Fisher | PA5-92037 | WB | 35 |
| p70(S6K) | Proteintech | 14485-1-AP | WB | 59 |
| Phospho-p70(S6K) | Proteintech | 28735-1-AP | WB | 59 |
| β-actin | Cell Signaling | 4967 | WB | 42 |
| Anti-rabbit IgG HRP-linked | Cell Signaling | 7074 | WB |  |

### MMP assay

MMP was examined by assessing TMRM (Thermo Fisher Scientific). After SL, cells where incubation with 20 nmol/L TMRM and Hochest stain 33258 (30 min, 37 °C) in the dark for 1 hour. The membrane of the FlexCell plates was then cut and transfered to 6-well plates. The membrane was washed twice tenderly with PBS and then approximately 500 µl culture medium was added on top of the membrane to maintain cell viability. A Leica Thunder Widefield fluorescence microscope was utilized for analysis.

### Live-Cell Imaging

Cellular morphology and MMP was assessed in real-time using the Incucyte^®^ S3 Live-Cell Analysis System (Sartorius, Ann Arbor, MI, USA). Cells were subjected to SL for 1 hour with the presence of 20 nM TMRM. Afterwards the FlexCell plate membrane was cut and transfered to 6-well plates and placed in the Incucyte^®^ System and the cell morphology and TMRM signaling were visualized continuously for 6 hours. The software was adjusted to obtain 9 images per well every 1 hour over the 6 hour period.

## Statistics

Data were analyzed using GraphPad Prism 7 (GraphPad Software, San Diego, CA) software. One-way analysis of variance (ANOVA) with Tukey’s multiple comparison (post hoc) test was performed in comparisons between more than two groups. Differences were considered statistically significant at a *p*-value of <0.05. All experiments were repeated successfully at least three times (i.e., at least three separate experiments were performed with cells isolated from different donors). All experimental samples were prepared in triplicates (*n* = 3).

## Data Availability Statement

The data that support the findings of this study are available in the methods, results of this article.

## Acknowledgments

We acknowledge the Biochemical Imaging Center at Umeå University and the National Microscopy Infrastructure, NMI (VR-RFI 2019-00217) for providing assistance in microscopy.

## Author Contributions

Conceptualization, Xin Zhou and Ludvig Backman; Data curation, Xin Zhou, Shaochun Zhu, and Junhong Li; Formal analysis, Xin Zhou, Junhong Li, Shaochun Zhu and Ludvig Backman; Funding acquisition, Ludvig Backman and Andre Mateus; Investigation, Xin Zhou, Junhong Li, Andre Mateus, Shaochun Zhu and Ludvig Backman; Methodology, Xin Zhou, Junhong Li, Shaochun Zhu and Ludvig Backman; Project administration, Andre Mateus and Ludvig Backman; Resources, Ludvig Backman and Andre Mateus; Supervision, Andre Mateus and Ludvig Backman; Validation, Andre Mateus and Ludvig Backman; Writing – original draft, Xin Zhou; Writing – review & editing, Ludvig Backman. All authors have read and agreed to the published version of the manuscript.

